# Single-cell imaging of protein dynamics of paralogs reveals mechanisms of gene retention

**DOI:** 10.1101/2023.11.23.568466

**Authors:** Rohan Dandage, Mikhail Papkov, Brittany M. Greco, Dmytro Fishman, Helena Friesen, Kyle Wang, Erin Styles, Oren Kraus, Benjamin Grys, Charles Boone, Brenda Andrews, Leopold Parts, Elena Kuzmin

## Abstract

Gene duplication is common across the tree of life, including yeast and humans, and contributes to genomic robustness. In this study, we examined changes in the subcellular localization and abundance of proteins in response to the deletion of their paralogs originating from the whole-genome duplication event, which is a largely unexplored mechanism of functional divergence. We performed a systematic single-cell imaging analysis of protein dynamics and screened subcellular redistribution of proteins, capturing their localization and abundance changes, providing insight into forces determining paralog retention. Paralogs showed dependency, whereby proteins required their paralog to maintain their native abundance or localization, more often than compensation. Network feature analysis suggested the importance of functional redundancy and rewiring of protein and genetic interactions underlying redistribution response of paralogs. Translation of non-canonical protein isoform emerged as a novel compensatory mechanism. This study provides new insights into paralog retention and evolutionary forces that shape genomes.

## Introduction

Gene duplication events play an important role in genome evolution and the emergence of adaptive complexity at the phenotypic level. Gene duplication occurs by polyploidy events resulting in whole genome duplicates (WGD) or duplication of small genomic regions, generating small-scale duplicates. During the divergence of paralogs, a large fraction of them is removed due to non-functionalization (Force et al. 1999). However, a significant proportion of the paralogs is retained, and this retention is thought to occur through three dominant routes (Kuzmin, Taylor, and Boone 2021): (1) by providing dosage amplification (Kondrashov and Kondrashov 2006) or backup compensation (Nowak et al. 1997) (2) by allowing subfunctionalization, i.e. the partitioning of functions between the sister paralogs (Force et al. 1999) and (3) by permitting neofunctionalization, i.e. the acquisition of novel functions (Ohno 1970; Casewell et al. 2011).

Systematic single gene deletion and loss-of-function screens in model organisms, such as yeast (Gu et al. 2003), worms (Conant and Wagner 2004), plants (Hanada et al. 2009), and human cells (De Kegel and Ryan 2019; Dandage and Landry 2019), have revealed that paralogs can provide genetic robustness. As a consequence, paralogs tend to have a lower fitness cost when perturbed in these screens when compared to the perturbation of singleton genes.

Genetic interactions have been used to study paralog divergence. Genetic interactions occur when a combination of mutations in different genes results in an unexpected phenotype, deviating from a model based on the integration of the individual mutant phenotypes (Costanzo et al. 2019). In yeast ∼30% of paralogs exhibit negative genetic interactions with each other, whereby the combined deletion of both paralogs results in a greater fitness defect compared to the effect of individual paralog deletions (VanderSluis et al. 2010). This finding indicates the pervasiveness of the compensatory functional redundancy between paralogs. Complex genetic interaction screens involving double mutants of dispensable WGD paralogs in yeast also revealed that functionally redundant paralogs tend to have a relatively large fraction of compensatory negative trigenic interactions (Kuzmin et al. 2020). According to the structural and functional entanglement model of duplicate divergence, the functional redundancy of paralogs is evolutionarily stable, as the divergence of functional domains is structurally constrained (Kuzmin et al. 2020). Thus, genetic interaction studies have uncovered the prevalence of functional redundancy among retained paralogs.

The mechanisms leading to paralog retention involving protein and RNA dynamics are not completely understood. Previous studies have uncovered evidence consistent with a “responsive backup circuit” model where inactivation of a paralog leads to the compensatory upregulation of a redundant sister paralog through changes to protein or mRNA expression levels (Kafri, Levy, and Pilpel 2006; DeLuna et al. 2010; Burga, Casanueva, and Lehner 2011). Other studies have shown changes in protein-protein interactions (PPIs) (Diss et al. 2017) and protein-DNA interactions (Gera et al. 2022) of paralogs. The function of a paralog can also be dependent on its paralog. In this scenario, termed dependency, the paralogs essentially act as a single functional unit and thus, in contrast to compensation model, inactivating a dependent paralog can lead to a fitness defect comparable to a singleton gene. Dependency is common among the heteromeric paralogs that physically interact with each other, as shown in yeast (Diss et al. 2017) and human cells (Dandage and Landry 2019).

While changes in mRNA expression, protein abundance, and protein interactions have been previously investigated as mechanisms of paralog retention, changes to protein localization have remained elusive despite evidence for subcellular localization differences among duplicated genes (Marques et al. 2008). The only known example of paralog compensatory localization changes comes from a previous study (Manolson et al. 1994) which investigated the localizations of Stv1 and Vph1 in the budding yeast. This paralog pair encodes proteins belonging to the V-ATPase protein complex, which carries out the acidification of organelles (Preston, Murphy, and Jones 1989; Manolson et al. 1992). These paralogs localize to distinct subcellular compartments: Stv1 resides in the Golgi and endosomes, whereas Vph1 is found in the vacuole. The growth defect caused by the *VPH1* deletion was rescued by the overexpression of *STV1*, and the protein it encodes relocalized to the vacuole suggesting compensation for the loss of *VPH1*. This example suggests that largely under-investigated relocalization-based protein dynamics of paralogs may play a fundamental role in compensation, and thereby in retention of paralogs.

In yeast, previous high-throughput microscopy studies (Tkach et al. 2012; Breker, Gymrek, and Schuldiner 2013; Chong et al. 2015; Kraus et al. 2017) reported relocalization of the proteome in response to environmental, chemical, or genetic perturbations and cataloged these response patterns (Lu et al. 2018). However, a systematic study of proteins through assessing their response to the loss of paralogs has been lacking. Such an investigation, afforded by the recent improvements in high throughput microscopy techniques and quantitative analytic approaches, could answer fundamental questions related to paralog retention such as: (1) at what prevalence proteins exhibit relocalization upon the deletion of their paralog, (2) how often the relocalizations are compensatory or dependent, (3) which properties of paralogs are predictive of relocalization, and (4) what mechanisms drive protein relocalization observed for paralogs.

In this study, we used a phenomics approach to capture subcellular distribution changes of proteins in response to their paralog deletions and identify the functional relationship of paralogs. We compared GFP-tagged strains of proteins to mutant strains harboring deletions of their paralogs, identified their subcellular relocalizations and quantified protein abundance changes. The correlative analyses of evolutionary and physiological gene features revealed that the subcellular distribution changes of proteins are associated with paralog functional redundancy. We uncover non-canonical protein isoform abundance, which refers protein variants of the canonical coding sequence, as a novel mechanism for paralog compensation.

## Results

### Quantifying paralog subcellular distributions using single-cell imaging

We constructed 328 yeast strains, enabling us to interrogate 82 WGD paralog pairs, each of which harbored a GFP-tagged protein in a wild-type or deletion of its paralog. These paralogs encoded proteins that exhibit distinct subcellular localizations (Chong et al. 2015; Koh et al. 2015; Breker, Gymrek, and Schuldiner 2013) and represent 82 of 551 of unique WGD paralog pairs (Byrne and Wolfe 2005). We used the Synthetic Genetic Array (SGA) approach to generate four strains per paralog pair by crossing strains carrying GFP-tagged proteins with strains carrying a wild-type or deletion of their paralog (Figure 1A, Methods). These strains enabled the assessment of the subcellular protein distribution changes in a reciprocal manner. Six proteins were paired with random paralogs to serve as negative controls. Using high-content microscopy, we measured changes to the subcellular localization and abundance of each protein in response to the deletion of its paralog (Figure 1B).

**Figure 1.**
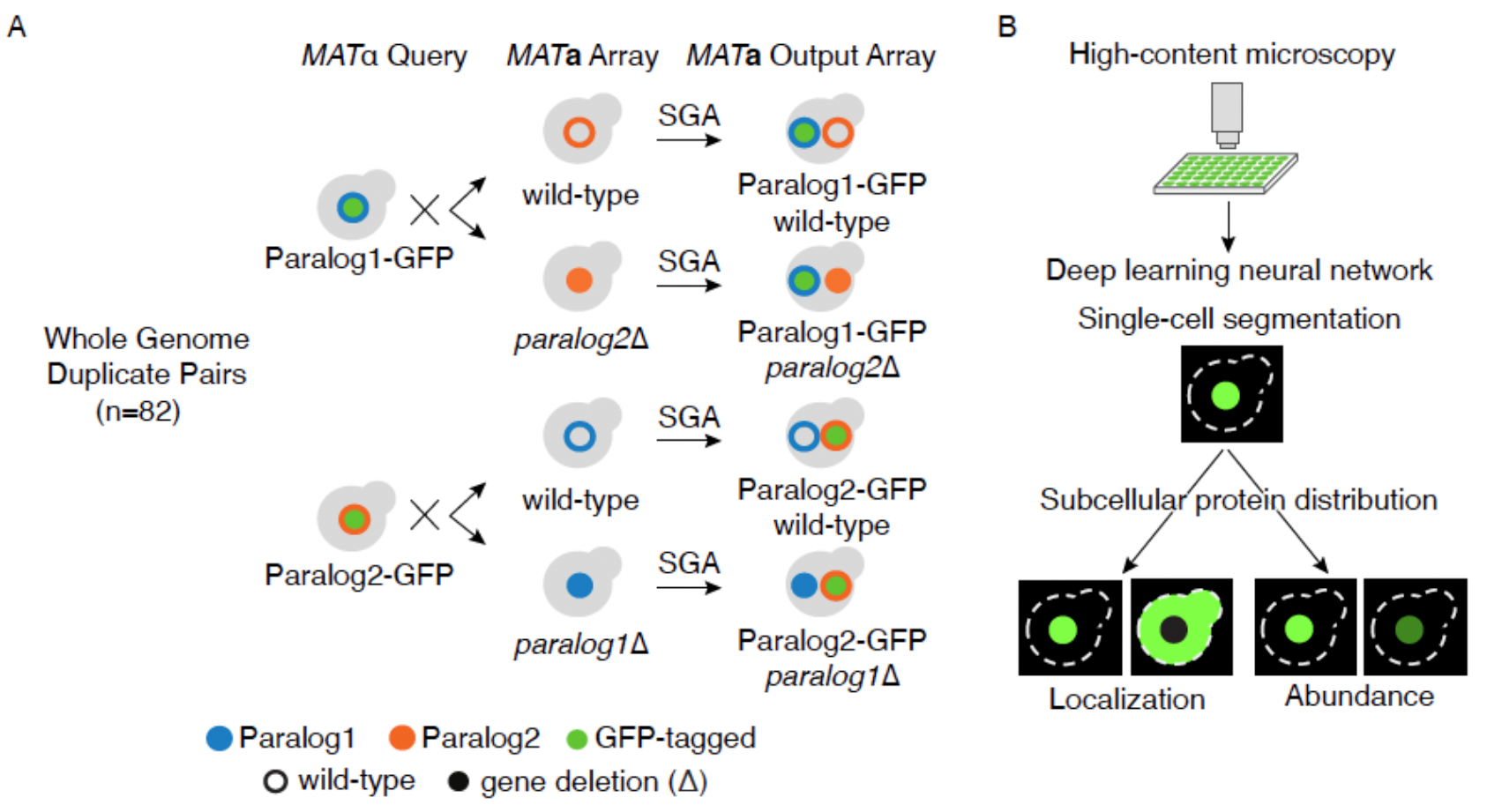
High-content protein dynamics analysis of paralogs. **A.** An illustration of strain construction for high-content screening of 82 paralog pairs originating from whole genome duplication (WGD) in yeast with previously reported localization differences. *MAT*α query strains harboring GFP-tagged proteins (green circle) were crossed with an array of *MAT***a** strains harboring either wild-type (open blue or orange circles) or deletion (Δ) (solid blue or orange circles) of their paralog. After induction of meiosis in heterozygous double mutants, sequential replica-pinning steps of the Synthetic Genetic Array (SGA) method were used to select the desired progeny, resulting in an output array of *MAT***a** strains harboring GFP-tagged proteins in wild-type and deletion backgrounds of their paralogs. **B.** High-content screening and analysis of the array of strains harboring GFP-tagged proteins in wild-type and deletion backgrounds of their respective paralogs. The strains were imaged using automated high-content microscopy. A deep neural network was used for cell segmentation and feature extraction to capture the subcellular protein distribution, which is a collective measure of protein abundance and localization.

We used an automated image analysis pipeline involving a deep neural network for single-cell segmentation and feature extraction. Images with artifacts or low number of cells were excluded from the analysis. Visual inspection of the resulting images revealed wild-type subcellular localization of paralogs that was consistent with previous studies (Chong et al. 2015; Huh et al. 2003). The neural network architecture, ResNet-18, was used to extract 128-dimensional features of single cells that were used to capture protein distributions, including protein localization and protein abundance (Data S1). We also obtained single-cell level protein abundances from the images by quantifying the mean pixel intensity per cell (Data S2). The resulting abundance scores in the wild-type background correlated well with previous measurements of protein abundance (Ho, Baryshnikova, and Brown 2018) (Figure S2A) and showed a strong correlation across replicates (Figure S2B).

### Protein dynamics reveal scenarios of compensation and dependency

To capture changes in subcellular localization and abundance of proteins in response to their paralog deletion, we calculated the changes in their protein distribution in the presence or absence of their paralog. We defined a redistribution score as the Euclidean distance between the centroid points of the extracted features comparing backgrounds for each protein (Figure 2A, Methods). The redistribution score is a collective measure of both the relative change in protein abundance and localization. We also separately quantified relative protein abundance changes for these paralogs.

**Figure 2.**
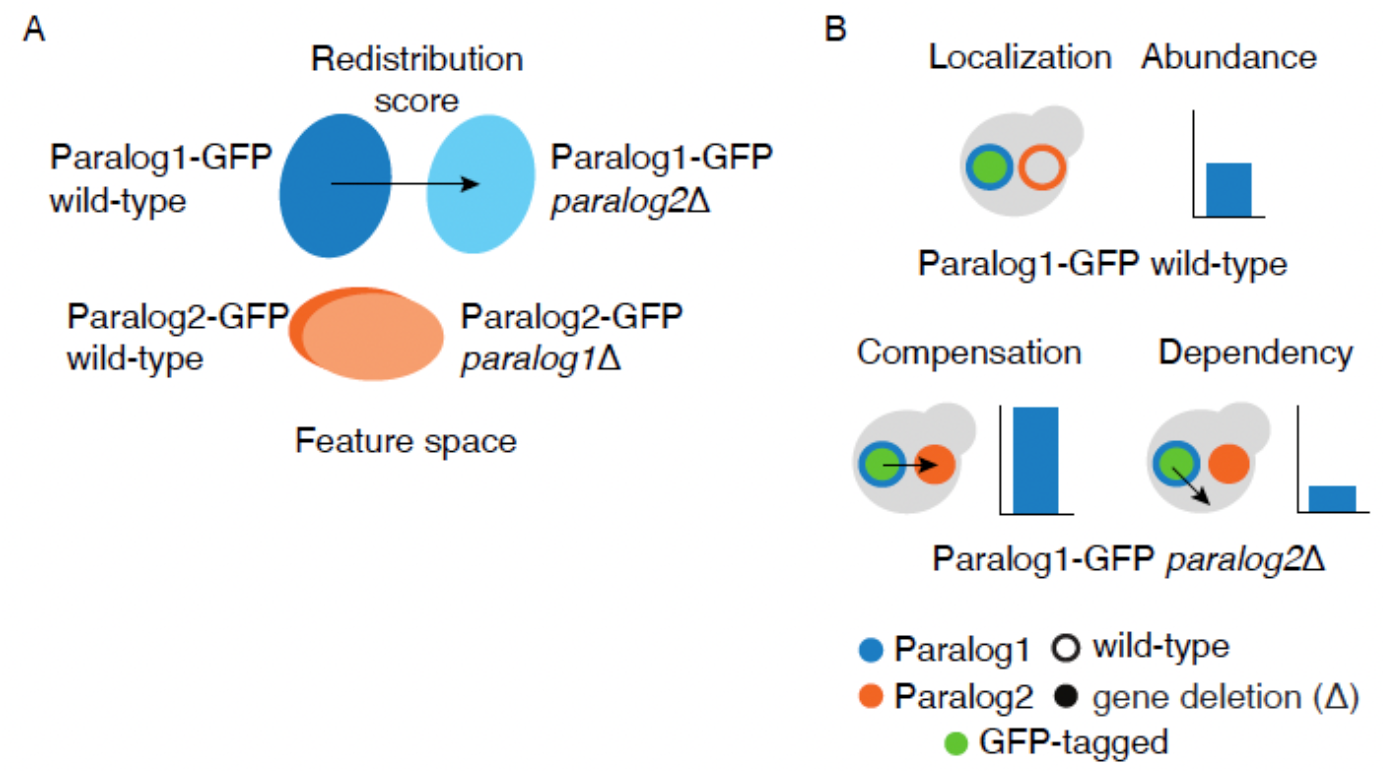
Identifying cases of compensation and dependency of paralogs using protein redistribution analysis. **A.** Schematic representation of the redistribution of the paralogs in the feature space (set of features extracted from deep neural network). The protein redistribution of a paralog between wild-type (dark blue/orange) and the deletion (light blue/orange) backgrounds is indicated by the arrow. **B.** Scenarios of the compensation and dependency for relocalization and relative abundance change. An example is depicted in which GFP-tagged protein 1 (green circle with blue border) is in the wild-type (open orange circle) or deletion (Δ) (solid orange circles) background of its paralog. Compensation refers to an increased protein abundance or relocalization to its paralogous protein subcellular compartment in response to the deletion of its paralog (left), whereas dependency refers to a decreased protein abundance or relocalization to a subcellular compartment distinct from its own or of its paralogous protein in response to the deletion of the paralog (right). Arrow denotes localization change.

The protein dynamics between paralogs should be captured by the response of proteins they encode to the deletion of their paralog when assessing their subcellular localization and abundance. Using the redistribution scores and relative protein abundance changes, we assessed the protein dynamics of paralogs. We predict that compensation would be observed when proteins respond to the loss of their paralog by changing their subcellular localization to reside in their paralog’s compartment and/or increasing their protein abundance. In contrast, dependency would be observed when proteins respond by diverging from their own or their paralogous protein wild-type subcellular localization and/or decreasing in abundance (Figure 2B).

### Proteins respond to their paralog loss by subcellular redistribution

In total, we quantified the redistribution scores for 164 proteins comprising 82 paralog pairs. These proteins showed a range of redistribution scores (Figure 3A), with 132 paralogs exhibiting a relatively low redistribution score (below 4.73), while a smaller subset of 32 paralogs belonging to 25 paralog pairs displayed a higher redistribution score (Figure 3B). The scoring was consistent with independent visual inspection, indicating that paralogs with relatively high redistribution scores exhibited subcellular localization and/or protein abundance changes, and randomly paired paralogs showed low redistribution scores (Table S2). This resulted in the classification of the redistributed and non-redistributed paralogs with minimal False Positive Rate (FPR) (Figure S3A).

**Figure 3.**
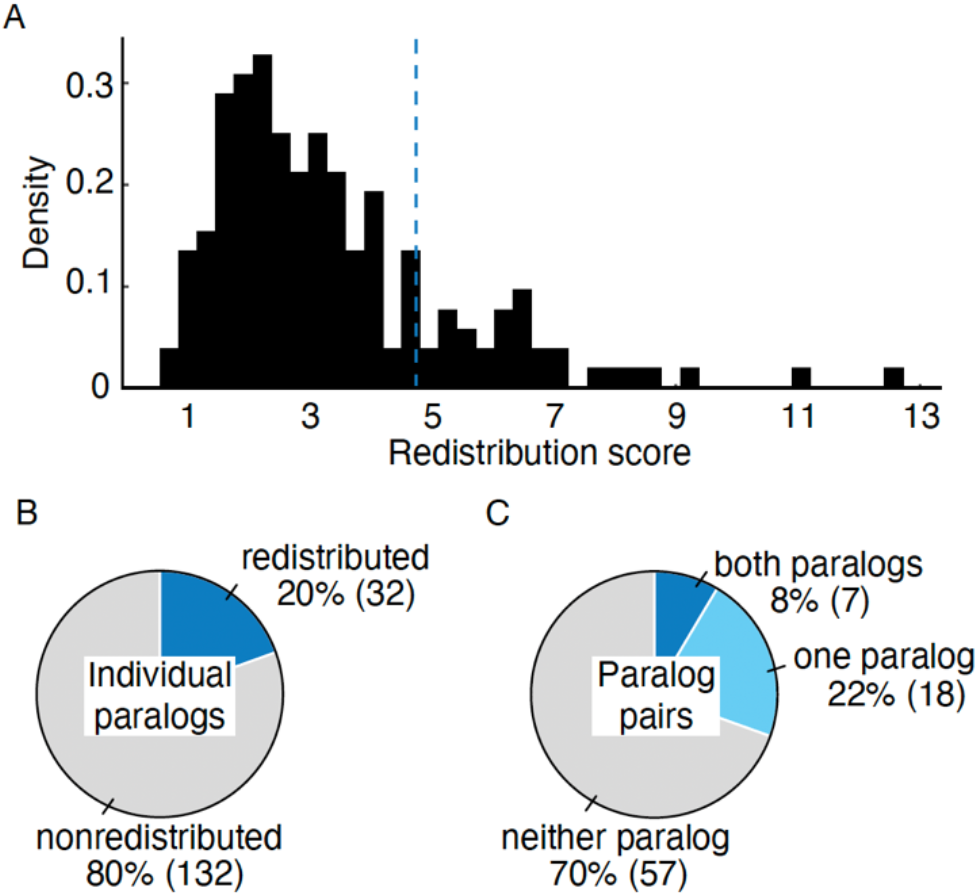
Paralogs respond to the loss of their sister paralog by protein redistribution. **A.** Histogram of the redistribution scores of screened paralogs. Redistribution score of greater than 4.73 (dashed line) was used to classify paralogs that exhibit protein redistribution. **B.** Pie chart showing the number of individual paralogs that exhibited redistribution in response to their sister paralog deletion (blue). Paralogs that show no response are depicted in grey. **C.** Pie chart showing the number of paralog pairs that exhibited redistribution in response to their sister paralog deletion. Pairs in which both paralogs responded to each other’s deletion are depicted in blue, those in which one paralog responded are in light blue, paralog that did not respond are in grey.

Paralog pairs having only one responsive member (18 pairs) were more common than reciprocal responsiveness, where both proteins respond to each other’s deletion (7 pairs, Figure 3C). This finding is consistent with the asymmetric divergence of paralogs (Kim and Yi 2006; VanderSluis et al. 2010).

### Proteins respond to paralog loss by abundance change

To test whether the redistribution score captures protein abundance changes, we separately quantified the abundance of each protein in the presence and deletion of its paralog (Figure 4A). We identified 30 proteins that exhibited a significant change in abundance upon the deletion of its paralog (|log_2_ fold change| ≥ 0.2, *p* < 0.05) (Methods). The relative protein abundance change is consistent with a previous study (DeLuna et al. 2010). For example, Cue4 (a protein of unknown function) and Pgm2 (a phosphoglucomutase which is involved in hexose metabolism (Boles et al. 1994)) were upregulated, whereas Sds24 (a protein involved in cell separation during budding (Jones et al. 2000)) was downregulated in both studies. Despite the differences in the methods used, the similarity in the classification supports the reproducibility of the relative abundance changes measured in the study. In total, 16 out of 30 proteins that showed significant relative abundance changes also exhibited redistribution. The remaining 14 proteins which were not classified as redistributed tended to show a modest protein abundance change (Figure S3B). Overall, there was an equal number of paralogs showing a relative increase in protein abundance, suggesting compensation, as there were paralogs showing a relative decrease in protein abundance, suggesting dependency (Figure 4B).

**Figure 4.**
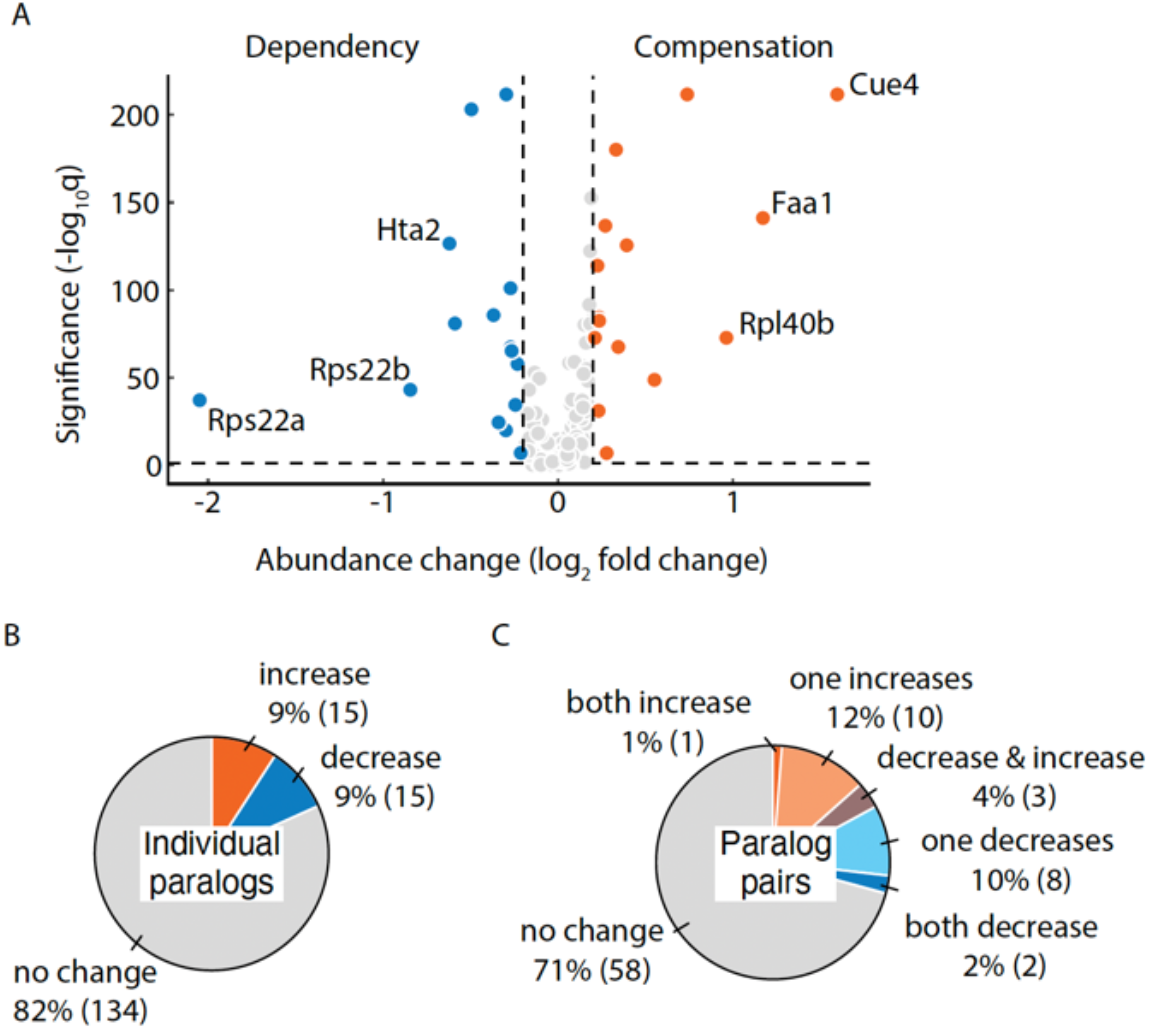
Proteins respond to their paralog loss by protein abundance changes. **A.** Volcano plot showing the protein abundance changes. The paralogs exhibiting significant relative abundance increase (log_2_ fold change ≥ 0.2, *q* < 0.05), suggesting compensation, are shown in orange, whereas the ones exhibiting decrease (log_2_ fold change ≤ -0.2 and *q* < 0.05), suggesting dependency, are shown in blue. The labeled points correspond to the top 3 proteins showing the greatest compensation and dependency. The points in gray correspond to the proteins that show non-significant changes. *q*: FDR-corrected *p*-value. **B.** Pie chart showing the number of individual proteins that exhibited significant protein abundance increase (orange), decrease (blue) or no significant change (grey) in response to their paralog deletion. **C.** Pie chart showing the number of paralog pairs that exhibited protein abundance change in response to their paralog deletion. Paralog pairs in which both proteins show increased abundance are shown in orange and those in which one protein increased are in light orange. Paralog pairs in which both proteins show decreased abundance are shown in blue and those in which one protein decreased are in light blue. Violet colour depicts the paralog pairs in which proteins responded in opposite directions. Paralog pairs that showed no response are shown in grey.

We also observed that compensation and dependency can co-occur in the same protein complex. Rps22a-Rps22b belong to the small ribosomal subunit and both show decreased abundance in response to each other’s loss indicating reciprocal dependency (Figure 4A, S4A-C). On the other hand, Rpl40a-Rpl40b belong to the large ribosomal subunit with Rpl40b increasing in abundance due to the loss of its paralog suggesting compensation (Figure 4A, S4D-F). Similar to redistribution (Figure 3C), the relative abundance changes in one member of the paralog pair were more common than reciprocal changes in both paralogs (Figure 4C). This observation was not confounded by the asymmetry in the native abundances of the paralogs. There was no significant association between the relative abundance changes and (1) the asymmetry (*p* = 1, Mann–Whitney U test) or (2) the under-expression of one of the paralogs (*p* = 0.6, Mann– Whitney U test) in the wild-type background. Therefore, the asymmetric relative abundance changes reflect the property of protein response to their paralog loss.

### Subcellular relocalizations of paralogs reveal cases of compensation and dependency

To further explore paralog redistribution, we considered subcellular localization changes. In total, among 32 proteins that showed redistribution in response to their paralog deletion, we used visual inspection and detected relocalization in 9 proteins (Cue4, Gga1, Hms2, Por2, Rpl40b, Rps22a, Rps22b, Upa1 and Upa2) from 7 paralog pairs (Figure 5A, S4, Table S4). Reciprocal cases in which both proteins relocalize are rare (2 of 7, Rps22a-Rps22b and Upa1-Upa2), similar to the rarity of reciprocal protein abundance changes. Relocalization was accompanied by protein abundance changes for 5 out of 9 proteins, indicating that redistribution captures both changes. The remaining 12 out of 32 proteins that were scored as redistributed were neither classified as relocalized nor showed relative abundance change. These paralogs may exhibit relocalization too subtle to be confidently detected by visual inspection but may still represent real biological response.

**Figure 5.**
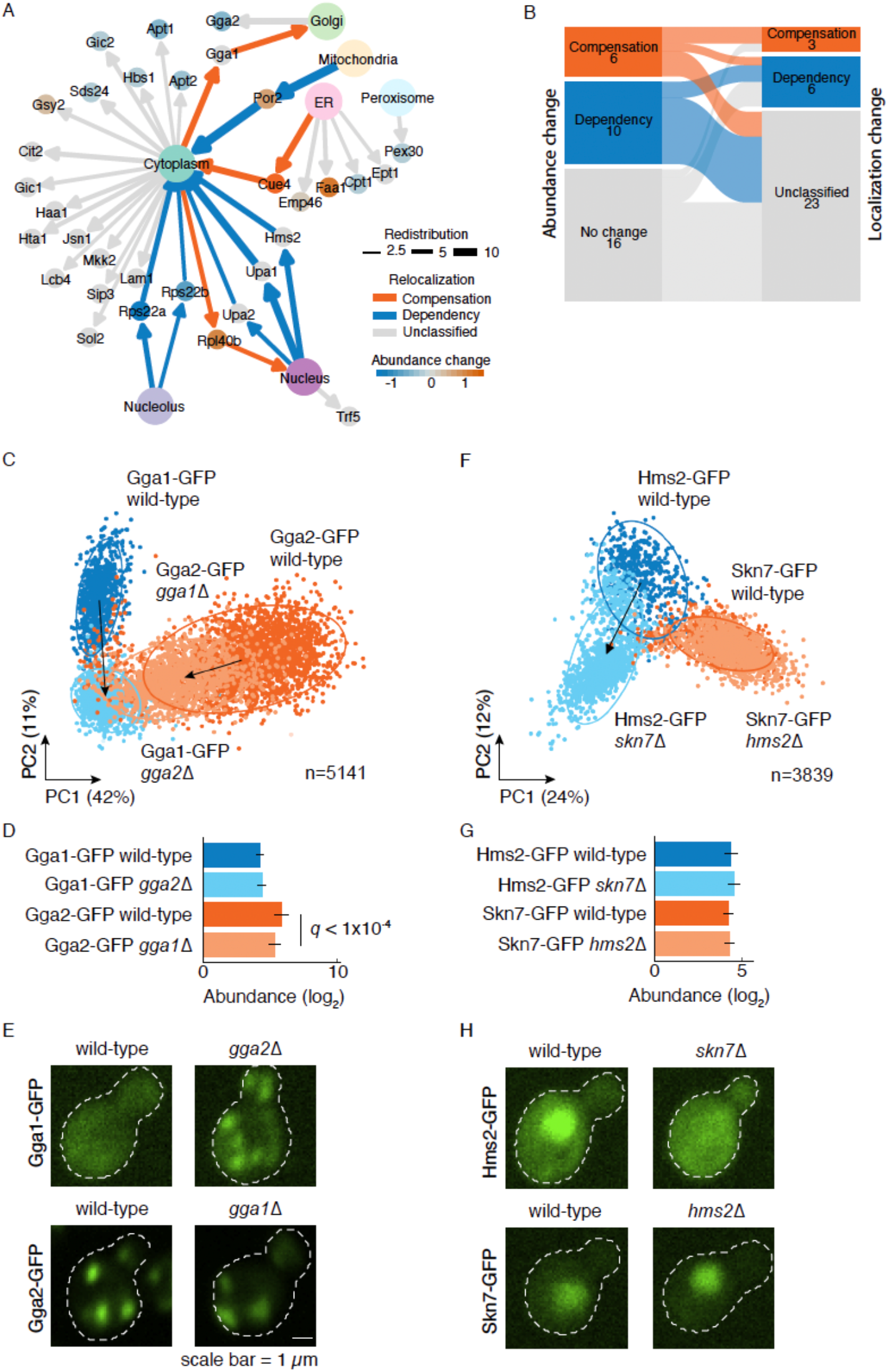
Proteins redistribute in response to their paralog loss by abundance and localization changes. **A.** Summary network of paralog redistribution. Redistribution score (edge width), abundance change (node color), protein relocalization (edges directed to subcellular compartment). Edges show relocalization and nodes show abundance change. The compensation and dependency are shown by the blue and orange colors, respectively. **B.** Comparison of proteins showing compensation (blue) and dependency (orange) between the relative abundance change and relocalization. **C.** Redistribution of the Gga1-Gga2 pair. Principal Component Analysis (PCA) of z-score normalized features is shown for each construct. Redistributed paralogs are shown with the arrow connecting the centroids of the clusters corresponding to the wild-type (dark blue/orange) and deletion (light blue/orange) backgrounds. The percentage variances explained by the PCs are indicated in parentheses. **D.** Relative abundance changes of the Gga1-Gga2 pair. The error bars show the 95% confidence intervals of the means. *q*: FDR-corrected *p*-value. Gga1-GFP is shown in blue and Gga2-GFP in orange. Wild-type paralog background is shown in dark blue/orange and deletion paralog background in light blue/orange backgrounds. **E.** Micrographs of representative yeast cells showing compensatory relocalization of Gga1-GFP in response to *GGA2* deletion and abundance dependency of Gga2-GFP in response to *GGA1* deletion. **F-H.** Similar to panels as **C**-**E**, but for the Hms2-Skn7 pair. The redistribution, relative abundance change, and micrographs are shown. The micrographs in panel **H** show the dependent relocalization of Hms2-GFP in response to *SKN7* deletion.

Considering proteins that respond to the deletion of their paralog by subcellular relocalization, we categorized three proteins as exhibiting compensation: Cue4, Gga1, and Rpl40b and six proteins as exhibiting dependency: Hms2, Por2, Rps22a, Rps22b, Upa1, and Upa2 (Figure 5A). We find that the compensation and dependency of relocalization and relative abundance changes often co-occur (Figure 5B). The redistributed proteins that show abundance compensation are more likely to show relocalization compensation, such as Cue4 and Rpl40b, except for Por2, which shows dependent relocalization. Similarly, redistributed proteins that show abundance dependency also exhibit relocalization dependency. Overall, among the redistributed paralogs, abundance and relocalization dependency was more prevalent than compensation.

Independent of the high-throughput screen, we re-imaged and validated two paralog pairs exhibiting redistribution, Gga1-Gga2, and Skn7-Hms2 (Methods), which represent cases of relocalization compensation and dependency. Among them, the Gga1-Gga2 pair is involved in facilitation of Golgi trafficking (Zhdankina et al. 2001). This pair is reciprocally redistributed (Figure 5C), which is due to Gga2 dependent relative abundance change (Figure 5D) and Gga1 compensatory relocalization from the cytoplasm to mitochondria, the wild-type localization of Gga2 (Figure 5E).

On the other hand, Skn7-Hms2 is a paralog pair that showed dependent localization. Skn7 is a transcription factor required for the optimal induction of heat-shock response to oxidative stress (Raitt et al. 2000) and osmoregulation (Janiak-Spens, Cook, and West 2005). Hms2 is similar to heat shock transcription factors, although poorly characterized. It redistributes in response to *SKN7* deletion (Figure 5F), with no abundance change (Figure 5G), but relocalizes from the nucleus to the cytoplasm (Figure 5H). The cellular compartment where Hms2 relocalizes is different from its wild-type localization and that of its sister, indicating a dependent relocalization. Overall, we found that the relocalization and the relative abundance changes of paralogs can collectively or individually manifest as their redistribution.

### Properties of paralogs are predictive of redistribution

After establishing the extent of redistribution and compensation, we next asked which paralog properties can predict these phenomena. Paralogs with a relatively high redistribution score are more likely to exhibit overlapping PPIs and have a shorter path length indicating that they are closely situated in the PPI network (Figure 6A). We find that paralogs with negative genetic interactions with each other were associated with a relatively high redistribution score (Figure 6B, left), indicating that the loss of both paralogs in a pair leads to a greater fitness defect compared to the fitness defect due to the loss of individual paralogs. A high redistribution score was also associated with a high trigenic fraction (Figure 6B, middle) due to a greater number of compensatory negative trigenic interactions compared to paralog-specific digenic interactions, indicating paralogs with a functional overlap. Also similar to PPI network, paralogs with a relatively high redistribution score were also closely located in the genetic interaction network (Figure 6B, right). Together these findings suggest that the functional redundancy may act as a prerequisite for paralogs to exhibit redistribution.

**Figure 6.**
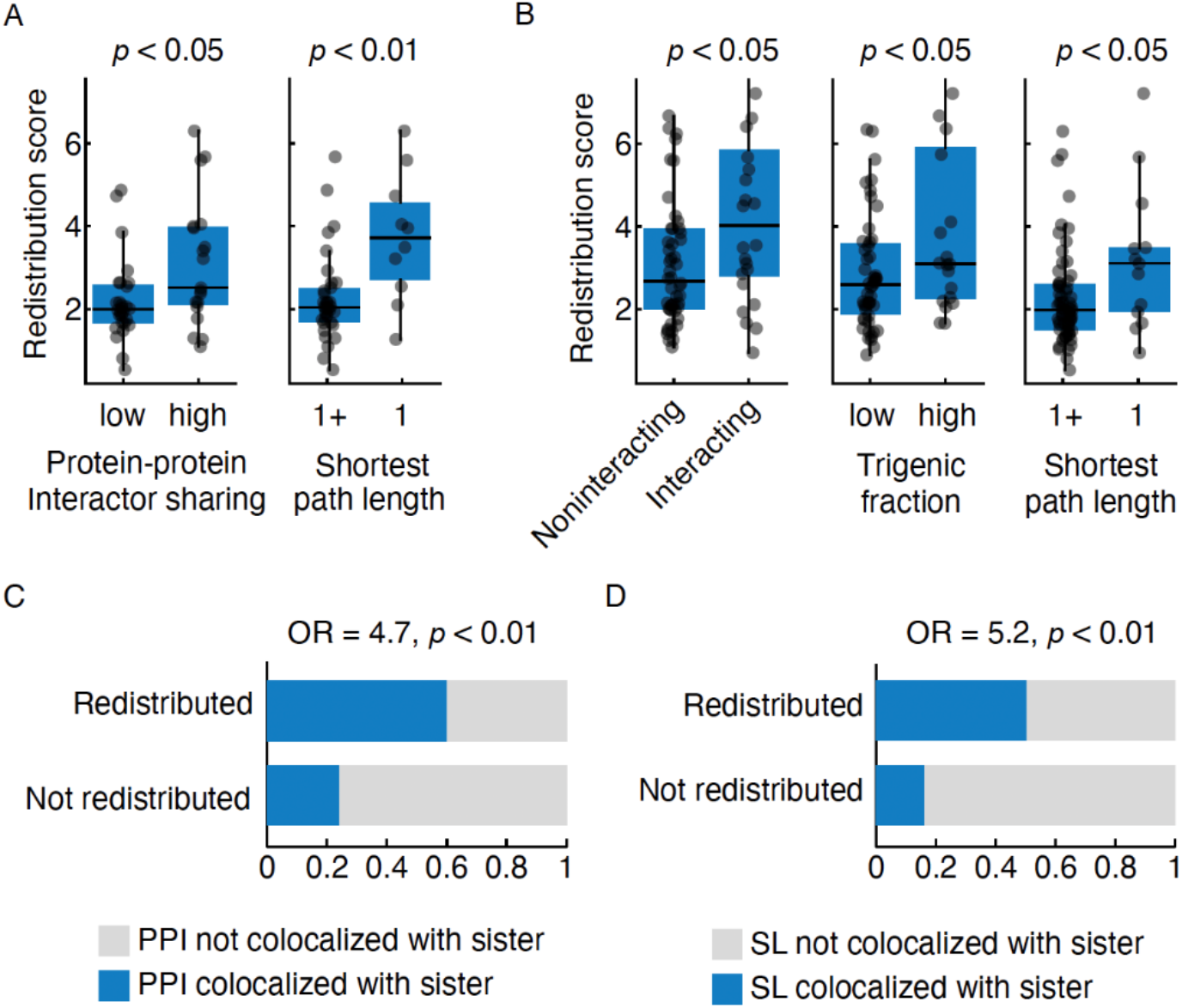
Redistribution correlates with protein-protein and genetic interaction network features of paralogs. **A.** Protein-protein interaction (PPI) network features. Plots show redistribution scores of individual paralogs, which were binned according to their shared PPI with their paralog (left) or the shortest path length between paralogs (right). Paralogs with fewer or more than the median number of shared interactors are denoted as low and high, respectively. The shortest path length of 1 (denoted as 1) and more than 1 (denoted as 1+) in the interaction network is shown. Merged PPI functional standard was used (Methods). **B.** Genetic interaction network features. Plots show redistribution scores of individual paralogs, which were binned according to the genetic interaction network features, including: (left) paralogs that exhibit a negative genetic interaction (interacting) or no interaction (noninteracting) within the paralog pair (previously defined stringent threshold i.e. *p* < 0.05, ε < -0.12) (Costanzo et al. 2016); (middle) paralogs belonging to the previously defined high and low trigenic interaction fraction classes (Kuzmin et al. 2020); and (right) the shortest path length of 1 (denoted as 1) and more than 1 (denoted as 1+) in the genetic interaction network (stringent threshold i.e. *p* < 0.05 and ε > 0.16 or ε < -0.12) (Costanzo et al. 2016). **C.** Redistributed proteins show interactions in the subcellular compartment of their paralog. The association of the redistribution and private interactors’ colocalization with paralog, is shown for PPI (left) and the synthetic lethal genetic interactors (right) (*p* < 0.05 and ε < -0.35) (Costanzo et al. 2016). OR: Odds ratio; *p*: p-value obtained from Fisher’s exact test. In the boxplots, the central line indicates the median, the extent of the box is from the first quartile (Q1) to the third quartile (Q3), and the whiskers extend to Q1-1.5*IQR and Q3+1.5*IQR. Statistical significance was determined using a two-sided Mann-Whitney U-test.

The subcellular redistribution of paralogs in response to the deletion of their sister paralogs may also be related to network rewiring. We found that compared to non-redistributed paralogous proteins, redistributed paralogous proteins are more likely to interact with other proteins residing in the subcellular compartment of their paralog (Figure 6C). This trend was also recapitulated using genetic interactions, whereby redistributed paralogs showed synthetic lethal interactions with proteins residing in the subcellular localization of their paralog (Figure 6D). We restricted this analysis to the private interactors, which are unique to each paralog, to eliminate the confounding effects of the shared interactors of the paralogs. This finding suggests that possessing interactors in the paralog localization might be a prerequisite for network rewiring associated with redistribution.

To gain finer insights into how interactions may change due to paralog loss, we compared the redistribution of paralogs with previously reported PPI changes of paralogs (Diss et al. 2017). Consistent with our expectation, among 10 paralogs that overlapped both studies, 7 paralogs with a PPI change exhibited relatively high redistribution scores (Figure S5). These findings suggest that PPI changes are predictive of redistribution. Thus, protein redistribution in response to their paralog loss occurs due to network rewiring which is likely mediated by compartment-specific PPIs.

### Compensatory relocalization and noncanonical protein isoform abundance

To further explore the mechanisms underlying paralog redistribution, we examined in detail the *CUE1-CUE4* paralog pair. The pair encodes proteins containing a Coupling Ubiquitin conjugation to ER degradation (CUE) domain that binds to ubiquitin (Shih et al. 2003). Cue1 is involved in the ER-associated degradation (ERAD) pathway for misfolded proteins (Bagola et al. 2013), whereas *CUE4* encodes a protein of unknown function. Cue1 is localized in the ER and cytoplasm, whereas Cue4 is predominantly localized in the ER (Huh et al. 2003; Koh et al. 2015). The Cue4 protein exhibited redistribution (Figure 7A), which was due to a compensatory relative abundance increase (Figure 7B) and compensatory relocalization to the cytoplasm (Figure 7C). This suggests an overall compensatory redistribution for the loss of *CUE1*.

**Figure 7.**
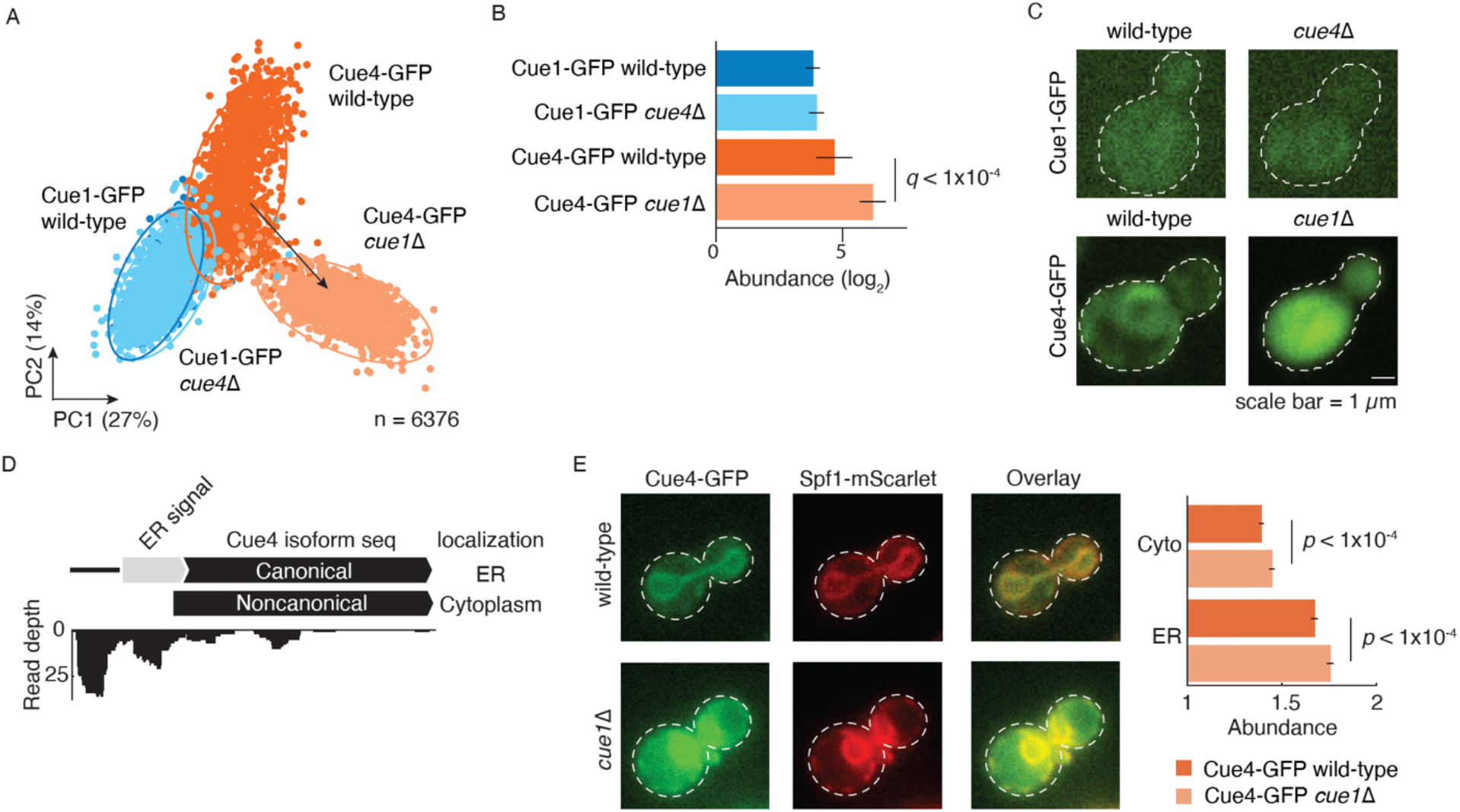
Redistribution of Cue4 and non-canonical protein isoform abundance. **A.** Redistribution of the Cue1-Cue4 pair. Principal component analysis (PCA) of z-score normalized features is shown for each construct. For the redistributed protein, the arrow connects the centroids of the clusters corresponding to the wild-type (dark blue/orange) and deletion (light blue/orange) backgrounds of the paralog. The percentage variances explained by the PCs are indicated in parentheses. **B.** Relative abundance changes of the Cue1-Cue4 pair. The error bars show the 95% confidence intervals of the means. *q*: FDR-corrected p-value. Cue1-GFP is shown in blue and Cue4-GFP is in orange. Wild-type background is shown in dark blue/orange and deletion background of the paralog in light blue/orange. **C.** Micrographs of representative yeast cells from the high-throughput screen showing compensatory relocalization of Cue4-GFP in response to *CUE1* deletion. **D.** The canonical and non-canonical protein isoforms of Cue4 (top) and corresponding read depths of ribosome footprints (Eisenberg et al. 2020) are shown. Predicted subcellular localizations of Cue4 isoforms are on the right. **E.** Cytoplasmic and ER abundance of Cue4-GFP in response to *CUE1* deletion was monitored. Spf1-mScarlet was used as a reporter of ER. The error bars show the 95% confidence intervals of the means, p-value calculated using Mann-Whitney U test.

Since Cue4 contains an N-terminal ER targeting signal sequence, we hypothesized that an N-terminal truncation might render the protein cytoplasmic. We mined the Translation Initiation Site (TIS) profiling data (Eisenberg et al. 2020) and identified a noncanonical isoform of Cue4. This isoform lacks the N-terminal ER targeting signal sequence due N-terminal truncation caused by the translation from an alternative start site. It is therefore predicted to be localized to the cytoplasm (Figure 7D). This prediction is robust to GFP-tagging of the protein sequences (Methods). A previously reported gain of an interaction between Cue4 and Ilm1, a poorly characterized cytoplasmic protein, in *cue1*Δ mutant background is consistent with the presence of a noncanonical cytoplasmic Cue4 protein isoform (Figure S5D) (Diss et al. 2017). Validation experiments in which we quantified Cue4 abundance in the ER and the cytoplasmic compartments showed that Cue4 protein abundance was greater in ER and cytoplasm in *cue1*Δ mutant compared to the wild-type background (Figure 7E). The Cue4 compensatory increase in abundance is consistent with the large-scale screen and the observed redistribution of Cue4 to the cytoplasm is due to increased translation of both protein isoforms.

## Discussion

Investigating protein dynamics underlying paralog retention has the potential to uncover how the fundamental process of genome evolution is fueled by gene duplication. In this study, we identified proteins that responded to the deletion of their WGD paralog by subcellular protein redistribution (Figure 3A), capturing relative abundance and localization changes (Figure 5A).

Protein-protein and genetic interaction network features revealed functional similarity as a predictor of paralog redistribution (Figure 6A&B). Functional similarity is considered a key characteristic of functional redundancy, which has been shown to be evolutionary stable (Li, Yuan, and Zhang 2010, Kuzmin et al 2020). In this study we show that functional redundancy plays an important role in subcellular protein dynamics of paralogs. We also find that rewiring of PPIs in specific subcellular compartments may guide the relocalization of paralogs (Figure 6C&D).

In our data, we observed that the paralogs from the small ribosomal subunit exhibited responses that were opposite to those of the paralog from the large subunit. This finding adds to the recent recognition that ribosomes are not homogenous, but rather heterogeneous macromolecular complexes (Genuth and Barna 2018). Ribosomal duplicate genes generate heterogeneity in the compositions of the ribosomes and carry out moonlighting functions (Komili et al. 2007; Segev and Gerst 2018), which collectively contribute to the robustness and phenotypic plasticity (Hu et al. 2022).

Our finding that Hms2 shows a dependent localization change in response to *SKN7* deletion is consistent with this pair thought to form a heterodimer (Gera et al. 2022). The asymmetric nature of the dependency is likely due to homomerization of Skn7 (Mariño-Ramírez and Hu 2002) but not of Hms2, which enables Skn7 to function in the absence of Hms2 but not vice versa. This is also consistent with a previous study examining DNA binding preferences, which showed that Hms2 loses binding to its target genes in response to the deletion of *SKN7* (Gera et al. 2022).

The analysis of alternate translation start sites of Cue4 is consistent with its compensatory relocalization occurring due to non-canonical protein isoform abundance. Similar to our findings in *cue1*Δ background, a previous study also reported that Cue4 relocalizes to the cytoplasm in the *rpd3*Δ background (Chong et al. 2015; Koh et al. 2015). Rpd3 is a lysine deacetylase with a diverse set of substrates (Kaluarachchi Duffy et al. 2012), and thus it is possible that it shares common substrates with ubiquitin ligases that interact with Cue1 and impinge on Cue4 function. The similar effect of the loss of *RPD3* and *CUE1* on Cue4 may reflect their roles in maintaining proteostasis, albeit in distinct ways. Rpd3 is required for the inactivation of ribosomal genes (Sandmeier et al. 2002); therefore, its deletion may lead to dysregulation of translation and proteostatic stress. Because Cue1 is involved in protein degradation (Bagola et al. 2013), its deletion may increase the load of undegraded proteins. Therefore, both *cue1*Δ and *rpd3*Δ might have a converging effect of increased proteostatic stress. Thus, Cue4 relocalization to the cytoplasm and its increased abundance could relieve the proteostatic stress by enabling additional protein degradation. Collectively, our study provides a novel molecular mechanism explaining subcellular protein dynamics of paralogs that underlie their retention within the yeast genome. The mechanisms of relocalization changes of paralogs were hypothesized (Diss et al. 2014) but never systematically tested prior to this study.

We found variation in protein dynamics, as captured by protein relocalization and abundance changes. Reciprocal redistribution of paralogs was found to be rare. We speculate that the asymmetric nature of the divergence of the paralogs may be the fundamental reason for the asymmetry. In cases where reciprocal redistribution did occur, both paralogs responded by compensation or both by dependency. This finding was consistent with a previous study examining PPIs of paralogs (Diss et al. 2017) and “monochromatic” genetic interactions involving members of pathways and protein complexes, indicating cellular organization into functional modules (Costanzo et al. 2010; Segrè et al. 2005).

In yeast, since WGD occurred around the time of the emergence of fruiting plants (Wolfe and Shields 1997), changes to the metabolic milieu may have exerted selective pressures for paralog retention. To account for such effects, carrying out the screening under different environmental conditions would be necessary to reveal environment-specific paralog responses. Such screenings would also reveal which protein dynamics are robust to environmental changes and which are not.

Extending the approach of phenomic screenings as presented in this study to human cells, where paralogs are more prevalent than in yeast, could reveal unexplored mechanisms explaining paralog retention and the genetic robustness of human cells (Dandage and Landry 2019). Such an extension may also help in characterizing potential subcellular relocalizations of human paralogs due to non-canonical isoforms resulting from splicing. The coupling of functional redundancy of paralogs with splice forms, which can be dubbed as “internal-paralogs” (Iñiguez and Hernández 2017), may be the key to explaining the exceptional genetic robustness of human cells.

Furthermore, such fundamental understanding could reveal novel cancer interventions, where synthetic lethality-based precision oncology therapeutic strategies could be devised to inhibit compensatory paralogs by blocking their redistribution, thereby specifically killing tumor cells while leaving normal cells unharmed.

The models of paralog retention, which have been outlined through decades of research in genomics and functional genomics, have paved the way to explore the underlying mechanisms of protein dynamics. Recent advances in phenomic screening and analysis approaches (Grys et al. 2017) are opening new avenues for increasing the scale of these screens and the resolution to the single-cell level. Our study demonstrates the potential of such technologies in addressing fundamental questions in the broad field of genome evolution.

## Methods

### Strain construction

#### Strain maintenance

All query strains were maintained on YEPD media (1% yeast extract, 2% peptone, 2% glucose, 0.012% adenine) supplemented with 100 μg/mL nourseothricin (clonNAT) (Werner Bioagents) and array strains on YEPD supplemented with 200 μg/mL geneticin (G418) (Agri-Bio). Single gene deletion strains were obtained from the SN query strain collection (Costanzo et al. 2016) and GFP strains were obtained from the GFP collection (Huh et al. 2003)

#### Query strain construction

We constructed and successfully screened 328 yeast query strains, each harboring a GFP fusion in wild-type or deletion background of its respective WGD paralog (Table S1). All duplicated genes originating from whole genome duplication (Byrne and Wolfe 2005) that were in the O’Shea collection (Huh et al. 2003), https://yeastgfp.yeastgenome.org) were examined. Paralog pairs were selected that exhibited different subcellular localizations or that were given an “ambiguous” annotation. Those that were “ambiguous” were examined in the CYCLoPs data (Chong et al. 2015) and those with distinct subcellular localization patterns were selected.

Strains from the GFP collection (Huh et al. 2003) with the following genotype were used: *MAT***a** paralog::GFP::HIS3MX6 *his3*Δ*1 leu2*Δ*0 met15*Δ*0 ura3*Δ*0.* To construct kanR GFP strains, they were marker switched from *HIS3* to *kanMX4* by PCR-mediated recombination strategy using high-efficiency LiAc transformation. p116 plasmid was used to PCR amplify *kanMX4.* This plasmid is ampicillin resistant and was cultured in 2YT + 120 μg/ml ampicillin. Primers that were used for *kanMX4* amplification were:

*MX4-F (5’ 3’): ACATGGAGGCCCAGAATACCCT*
*MX4-R (5’ 3’): CAGTATAGCGACCAGCATTCAC*

Marker switch was confirmed by replica plating on SD-his to confirm His^-^ phenotype of the kanR transformants. The resulting kanR GFP strains were introduced into the desired deletion background SN# (*MAT**α** paralog*Δ*::natMX4 can1*Δ*::STE2pr-Sp_his5 lyp1*Δ *his3*Δ*1 leu2*Δ*0 ura3*Δ*0 met15*Δ*0 LYS2+*) or the control strain Y8835 (*MAT**α** ura3*Δ*::natMX4 can1*Δ*::STE2pr-Sp_his5 lyp1*Δ *his3*Δ*1 leu2*Δ*0 ura3*Δ*0 met15*Δ*0 LYS2+*) by SGA (Kuzmin et al. 2014, 2016a, 2016b). Three control pairs were included in which random paralogs were paired together.

Briefly, the strains were crossed, sporulated and *MAT***a** meiotic progeny were selectively germinated using *STE2pr-Sp_his5* marker.

Final strains were of the following genotypes (Table S1):

1. *PAR1::GFP::kanMX4 ura3Δ::natMX4 can1Δ::STE2pr-Sp_his5 lyp1Δ his3Δ1 leu2Δ0 ura3Δ0 met15Δ0 LYS2+*
2. *PAR1::GFP::kanMX4 par2Δ::natMX4 can1Δ::STE2pr-Sp_his5 lyp1Δ his3Δ1 leu2Δ0 ura3Δ0 met15Δ0 LYS2+*
3. *PAR2::GFP::kanMX4 ura3Δ::natMX4 can1Δ::STE2pr-Sp_his5 lyp1Δ his3Δ1 leu2Δ0 ura3Δ0 met15Δ0 LYS2+*
4. *PAR2::GFP::kanMX4 par1Δ::natMX4 can1Δ::STE2pr-Sp_his5 lyp1Δ his3Δ1 leu2Δ0 ura3Δ0 met15Δ0 LYS2+*

#### Microscopy

Cells were prepared for imaging and imaged as described in detail previously (Kuzmin et al. 2014; Cox et al. 2016). Briefly, cells were grown to saturation in 200 ml SDmsg medium (0.1% monosodium glutamate, 0.17% yeast nitrogen base without amino acids and without ammonium sulfate, 2% glucose, 0.15 g/l methionine) containing 200 μg/ml G418 and 100 μg/ml clonNAT in 96-well beaded microplates, then diluted in 800 ml SDmsg low fluorescence medium (with 0.17% yeast nitrogen base without amino acids and without ammonium sulfate and without riboflavin and folic acid) containing 200 μg/ml G418 and 100 μg/ml clonNAT in beaded deep-well blocks and grown overnight to early log phase. All strains were grown at 30°C. Cells at ∼0.2-0.4 OD600/ml were transferred to a 384-well Perkin-Elmer Ultra imaging plate and left to settle for ten minutes before imaging. Four images per well, each containing fifty to a hundred cells, were taken in a single plane using an automated spinning disk confocal microscope (Evotec Opera, PerkinElmer) with a 60x water-immersion objective.

## Image analysis

### Segmentation

#### 1. Annotation processing

We processed ground truth object annotation to provide a segmentation neural network with the right objective to efficiently separate individual cells in clumps. To transform object segmentation into binary data and distinguish between the individual stencils, we eroded each stencil with the minimal cross, i.e., we claim the pixel to be background if any of its four neighbors is from the background. After erosion, we performed a Euclidean distance transformation of the binary segmentation by labeling each pixel with the distance to the nearest background pixel using SciPy (Virtanen et al. 2020). The transformed result was clipped at 20 and scaled between 0 and 1. Finally, we took a weighted average between binary segmentation and the distance map with a coefficient of 0.8, i.e., border pixels of distance-transformed stencils obtained a value of 0.8, which gradually increased to 1 maximum towards the object center (Figure S1). Such target transformation forced the segmentation algorithm to focus primarily on object centers and effectively separate the cells.

#### 2. Image processing

Before further transformations, we calculated a reference background image and subtracted it from all the images in the respective dataset. To do so, noise in each coordinate was calculated as a median pixel value by this coordinate across all the images in the dataset. This was done to mitigate an uneven illumination pattern as well as the read-out noise across the microscope’s field of view.

Protein abundance varies greatly depending on the protein localization and query gene. Thus, the detected signal range in images obstructs segmentation networks from generalization and efficient convergence. To mitigate this problem, we applied a logarithmic transformation to the base of *e* to all the images and standardized the pixel values in each image to mean 0 and standard deviation 1.

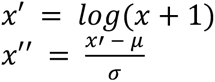

#### 3. Neural network architecture and training

For segmentation, we trained a fully convolutional U-Net neural network architecture (Ronneberger, Fischer, and Brox 2015) from the Segmentation models PyTorch library (Iakubovskii 2019). The training set was obtained from a previously manually annotated images of yeast cells that were imaged on the same microscope (available upon request). We utilized ResNet-34 (He et al. 2016) backbone as a network encoder. It consists of a single convolutional layer with 64 filters 7 x 7, followed by 16 residual units in blocks of 3, 4, 6, and 3 with 64, 128, 256, and 512 filters 3 x 3 respectively. Each residual unit consists of a layer sequence BN-ReLU-Conv-BN-ReLU-Conv with additively connected input and output. The first convolutional layer and each group of residual blocks are followed by a max-pooling layer and are considered to be a U-Net level. The encoder is followed by the decoder. The decoder consists of five levels Conv-ReLU-Conv-ReLU with 64 filters 3 x 3 per convolutional layer. Each encoder level is connected to the respective decoder level by a skip-connection. The final decoder layer outputs a single-channel probability map through the sigmoid activation function.

We implemented the network with PyTorch deep learning library (Paszke et al. 2019) in Python. Network parameters were initialized as proposed previously (He et al. 2015). We used the transformed distance maps (see Annotation processing) as training targets and optimized binary cross-entropy loss for the network’s predictions. The network was trained in the PyTorch Lightning framework (Falcon et al. 2020) for 200 epochs with a batch size of 32. Adam optimization (Kingma and Ba 2014) with default hyperparameters was used. We utilized the cosine annealing learning rate schedule (Paszke et al. 2019; Loshchilov and Hutter 2016) from an initial value 3 x 10^-4^ to 1 x 10^-7^.

For each image in a training batch, we randomly cropped patches 256 x 256 pixels from the original images and augmented them with random flipping and rotation by {0, 90, 180, 270} degrees. Additionally, each image was scaled and shifted by factors, sampled uniformly from ranges [0.5, 2) and [-1, 1) respectively. For the inference, we padded single full-size images with reflection until divisibility by 32 to adhere to network architecture requirements; outputs in padded areas were not analyzed.

We trained the network using 2246 images and validated its performance with 250 images after each epoch, using 1 validation image per gene label. We saved the best weights by a validation cross-entropy loss and used them in our following experiments.

#### 4. Post-processing algorithm

Individual cells are not distinguished perfectly in the segmentation network’s output. To split the clumps, we used the Watershed algorithm from the scikit-image library (van der Walt et al. 2014). The network predictions thresholded at 0.8 were used as watershed seeds, and a threshold of 0.5 was used to obtain object boundaries. After the watershed, we removed any objects smaller than 256 and larger than 8192 pixels by area.

### Feature extraction

We considered each object in the output segmentation map to be a cell. From each full-size image, we extracted all detected cell frames of size 64 x 64 pixels. We then trained a neural network classifier to distinguish cell frames between 182 gene classes in the wild-type context. Many of the genes share a subcellular localization, so we did not utilize the last dense classification layer of the network after the training. Instead, for each cell frame, we extracted 128 features from the penultimate pooling layer. These features do not directly map to the perceived cell properties, such as intensity, size, etc. However, they should provide sufficiently compressed information about the protein subcellular localization.

### Neural network architecture and training

For feature extraction, we trained a ResNet-18 neural network architecture. It consists of a single convolutional layer with 16 filters 3 x 3, followed by 4 residual blocks of 2 residual units each with 16, 32, 64, and 128 filters 3 x 3, respectively. Each residual unit consists of a layer sequence BN-ReLU-Conv-BN-ReLU-Conv with additively connected input and output. Feature extractor is concluded by a global average pooling layer, which yields 128 features for each input. During the training, these features were passed to the dense classification layer with 182 outputs followed by a softmax activation function. We used simple image augmentations from the Albumentations library (Buslaev et al. 2020): random horizontal and vertical flipping, shifting, scaling, and rotation with default parameters.

We implemented the network with PyTorch deep learning library (Paszke et al. 2019) in Python. Network parameters were initialized as proposed previously (He et al. 2015). We used encoded gene labels as training targets and optimized categorical cross-entropy loss for the network’s predictions. The network was trained in the PyTorch Lightning framework (Falcon et al. 2020) for 200 epochs with a batch size of 512. Adam optimization (Kingma and Ba 2014) with default hyperparameters was used. We utilized a previously defined the cosine annealing learning rate schedule.

We used all the wild-type cell frames from Field 4 for validation (∼25% of the total number) and the rest of the frames from Fields 1-3 for training. We did not distinguish between replicates at this stage.

### Filtering of the imaging data

The genes whose localization was mismatched with the known localizations (Chong et al. 2015), images containing abnormalities, artifacts, and excessively high heterogeneity, and the replicates containing very low cell numbers were removed from the dataset. In total, 8 paralog pairs were removed from the dataset based on this filtering procedure, resulting in 82 paralog pairs in the final dataset.

### Redistribution analysis

To detect differences of the subcellular protein distributions in the presence or absence of its sister paralog, we calculated Euclidean distances between the centroid points of the extracted features. We aggregated the 128 features for all cells from each strain in 3 replicates (mean number of cells ∼500) by calculating centroid points, which represent subcellular protein distributions. We refer to the distances between the centroid points for the wild type and deletion backgrounds as “redistribution scores”. For the visualization of the redistribution, we z-score normalized the features, and transformed them with the principal component analysis.

The threshold for classifying the paralogs based on the redistribution score was calculated using AUC-ROC analysis (Figure S3A). In this analysis, 12 visually inspected paralogs that showed relocalization and/or relative abundance change were used as true cases, and negative controls i.e. the six paralogs that were paired randomly were used as the false cases. The threshold providing the minimal False Positive Rate (FPR) was considered.

### Protein abundance quantification

We quantified the protein abundance by measuring raw pixel intensities in cell frames. For both conditions, we measured mean raw pixel intensity across all the pixels in all the frames in the condition, which we refer to as the abundance score.

For calculating the relative abundance changes, we first log-transformed (base = 2) the abundance scores with a pseudocount of 1 and then calculated the difference between the WT and deletion backgrounds. The statistical significance of the differences were calculated using two-sided Mann-Whitney U test, and corrected for multiple testing using Benjamini/Hochberg method.

### Experimental validation of relocalization

Strains of two redistributed paralog pairs (s*GGA1-GGA2* and *HMS2-SKN7*) representing cases of compensation and dependency were re-imaged to ensure reproducibility using Leica Confocal Microscope, 40X.

### Paralog features

#### Protein-protein interactions

PPI data were downloaded from BioGRID (Oughtred et al. 2021) (downloaded on 2022-06-27). The data was filtered to keep only physical ‘Experimental System Type’s and interactions from high-throughput studies (Krogan et al. 2006; Tarassov et al. 2008; Yu et al. 2008; Babu et al. 2012; Gavin et al. 2006).

#### Genetic interactions

Digenic interaction scores were obtained from a previous study (Costanzo et al. 2016). The subsets of the interactions i.e. lenient (*p* < 0.05), intermediate (*p* < 0.05 and interaction score |ε| > 0.08), and stringent (*p* < 0.05 and ε > 0.16 or ε < -0.12) were obtained by applying the previously defined thresholds. Trigenic interaction fraction classes were obtained from a previous study (Kuzmin et al. 2020).

#### Network features

The degree and the shortest path lengths were calculated using the networkx Python package (Hagberg, Schult, and Swart 2008).

#### Colocalization

The colocalization was classified based on the Jaccard index

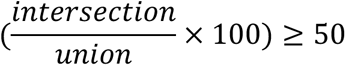

### Prediction of the relocalization of the non-canonical protein isoforms

The sequence of the non-canonical protein isoform of Cue4 protein was obtained from a previous study (Eisenberg et al. 2020). (Thumuluri et al. 2022)The Translational Initiation Site (TIS) profiling data for the vegetative exponential phase (SRR11777267) was used to visualize the read depth along the length of Cue4 in Figure 7D. Using the protein sequences of Cue4 and its non-canonical protein isoform, their localizations were predicted using DeepLoc2.0 (Thumuluri et al. 2022).

### Validation of Cue4-GFP redistribution

The strain Y109 *(MAT***a** *CUE4::GFP::kanMX4 cue1Δ::natMX4)* was reconstructed by PCR mediated-deletion, which replaced *CUE1* with a *natMX4* cassette using *MAT***a** *CUE4::GFP::kanMX4* starting strain. *CUE1* gene deletion was confirmed using colony PCR. Both Y109 and LocRed-6 (*MAT*a C*UE4::GFP::kanMX4 ura3Δ::natMX4)* strains were crossed to the strain Y110 harboring *SPF1-mScarlet,* an ER reporter (*MAT*α *SPF1-mScarlet-HYG*). Diploid strains were sporulated and tetrad dissected to select for the genotype of interest (NAT^R^, G418^R^, HYG^R^). The resulting strains (Y122 and Y123) were grown in YEPD to mid-log phase at 30°C. Live cells were rinsed once with water and once with low-fluorescent media (Sheff and Thorn 2004). Then the strains were imaged in low-fluorescent media using the Nikon Eclipse TiE inverted C2 confocal microscope at 100X oil immersion objective with filters for GFP (488/10x + 525/50m) and mScarlet (561/10x + 630/75m). DIC, GFP, and mScarlet were imaged sequentially. Refer to Table S1 for complete strain genotype details.

The individual channels from the raw images (Nikon NIS-Elements ND2 format) were separated using nd2 Python package (https://pypi.org/project/nd2). The images from the channel with the GFP intensity were used to carry out the cell segmentation, using YeastSpotter (Lu et al. 2019). The single cells were labeled using scikit-image (van der Walt et al. 2014) to assign a unique identifier to each cell. On average 50 cells per strain were quantified in three independent replicates. The cells at the edges of the images, within the median major axis length of the cells, were removed. The raw protein abundance per cell was calculated as the median value of the intensity, which was then divided by the intensity of the background to calculate the normalized abundance per cell, using htsimaging Python package (Dandage 2023a). Within each replicate, the images with extreme background intensity were discarded using the mean+/-stdev as thresholds. The images from the red channel containing the intensity for the ER reporter, Spf1-mScarlet, were used to identify the locations of the ER within the cells. Within each cell, the pixels with intensity ≥ 0.975 quantile were marked as the locations of the ER, whereas the ones with intensity < 0.973 quantile were marked as cytoplasmic. The cells with less than 100 ER pixels were discarded. Similar to the calculation of the cell-wise protein abundance, the protein abundances at the ER and cytoplasmic locations were calculated as the median value of the intensity at these locations divided by the background intensity of the image (Dandage 2023a).

Imaging data were analyzed using open-source Python packages such as scikit-image (van der Walt et al. 2014) and htsimaging (Dandage 2023a). The statistical analysis was carried out using scipy (Virtanen et al. 2020), statsmodels (Seabold and Perktold 2010), scikit-learn (Pedregosa et al., n.d.), numpy (Harris et al. 2020) and pandas (The pandas development team 2023). The network analysis was carried out using networkx (Hagberg, Schult, and Swart 2008). The data visualization was done using matplotlib (Hunter 2007) and seaborn (Waskom 2021). The Python package roux (Dandage 2023b) was used for both data analysis and visualization. The versions of these and other tools used in the analysis are provided along with the source code.

## Acknowledgments

We thank Malcolm Whiteway and Kuzmin lab members for discussions and critical comments on the manuscript. We thank Chris Law and the Center for Microscopy and Cellular Imaging at Concordia University for guidance on microscopy.

## Funding

Supported by Canada Research Chair program grant (CRC2021-00031 to E.K.), Canada Foundation for Innovation grant (CFI 41563 to E.K.), Natural Sciences and Engineering Research Council grant (RGPIN-2022-04674 to E.K.), Wellcome Trust (220540/Z/20/A to LP) and Revvity Inc. (VLTAT19682 to D.F. and L.P.), the National Institutes of Health (R01HG005853 to B.J.A., C.B.), and the Canadian Institutes of Health Research (PJT-180259 to B.J.A.). Computing resources and data storage services were partially provided by the High Performance Computing Center of the University of Tartu and the Digital Research Alliance of Canada.

## Author contributions

Conceptualization: R.D. and E.K.; Methodology and investigation: R.D., M.P., B.M.G., D.F., H.F., E.S., and E.K.; Formal analysis: R.D., M.P., B.M.G., D.F., C.B., B.J.A., L.P. and E.K.; Resources: K.W., O.K. and B.M.G.; Writing - original draft: R.D. and E.K.; Writing - review and editing: R.D., M.P., B.M.G., D.F., E.S., H.F, C.B., B.J.A., L.P. and E.K.; Supervision: E.K.; Funding acquisition: C.B., B.J.A., L.P. and E.K.

## Supplementary figures

**Figure S1.**
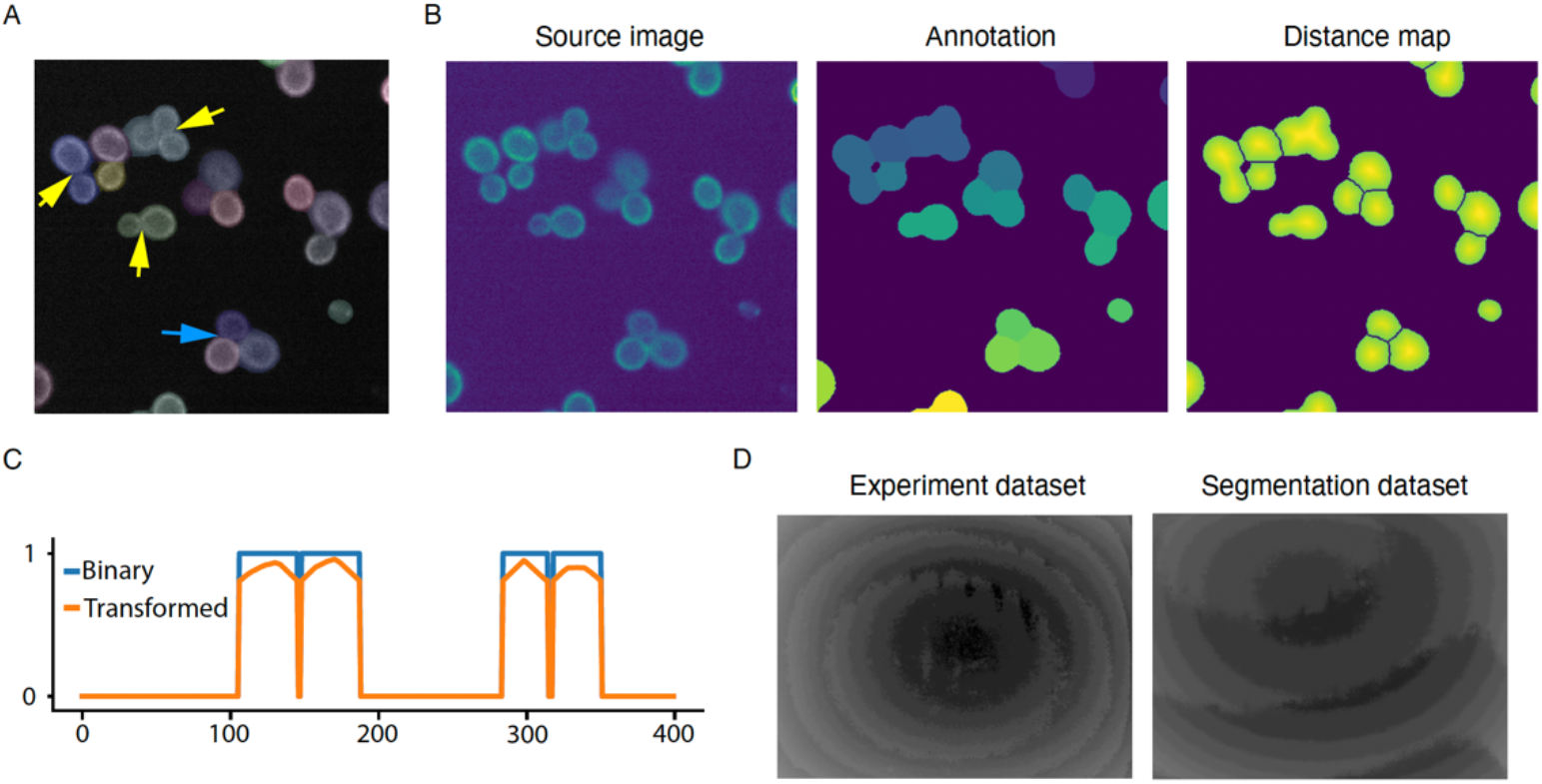
Optimization of the image segmentation. **A.** A representative cropped microscopy image, with object annotation overlayed. The yeast cells in the ground truth are shown using yellow arrows. An example of fully separated cells is shown using the blue arrow. **B.** Illustration of the proposed annotation processing into distance maps. Left: cropped microscopy image; center: ground truth object annotation; right: output distance map. **C.** Vertical cross-sections of the binary segmentation and the distance map. **D.** Log-transformed read-out noise highlights the uneven illumination pattern in both datasets.

**Figure S2.**
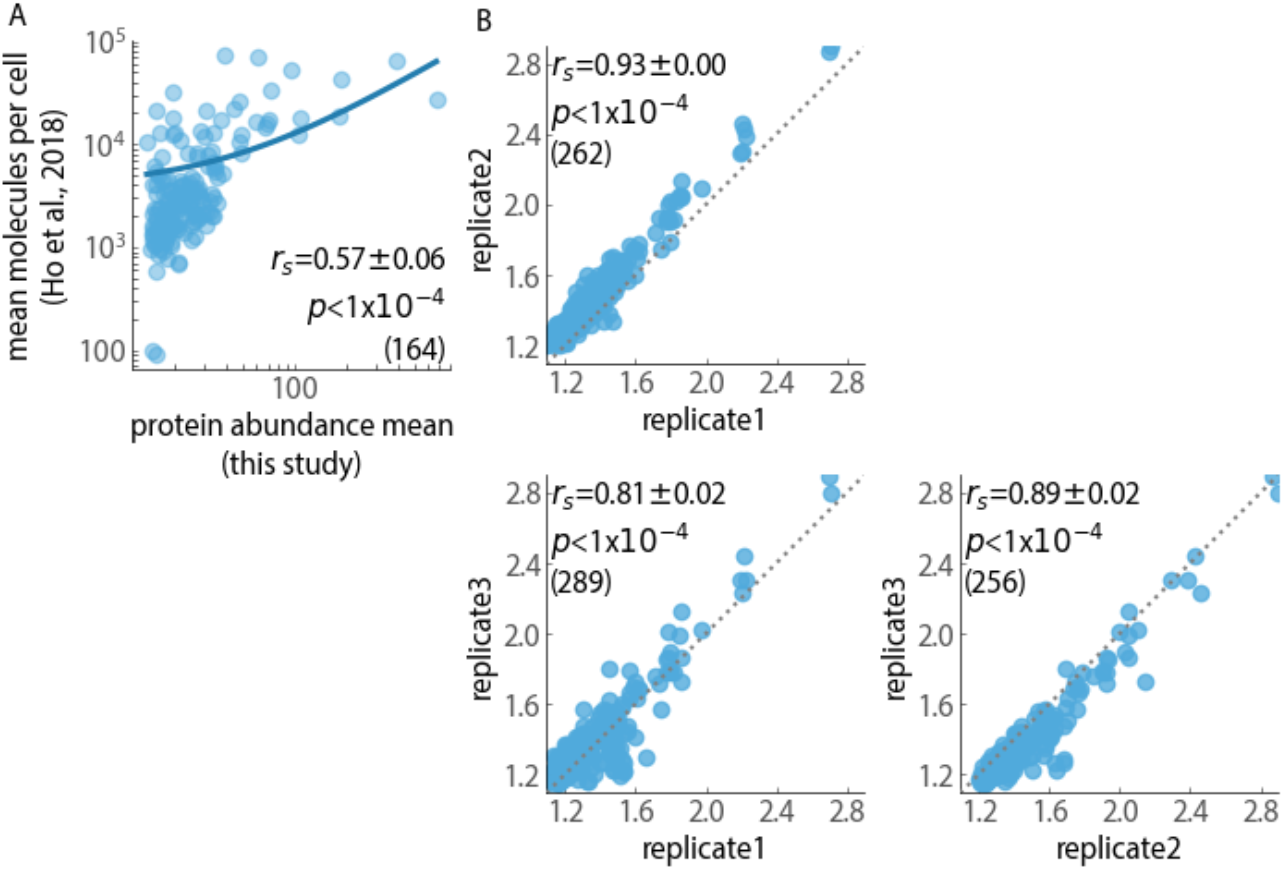
Protein abundance conformity with the known standard and reproducibility across replicate measurements. **A.** Correlation of the abundance values in the wild-type background with the reference values (Ho, Baryshnikova, and Brown 2018). The line indicates the fitted regression model of order 1. **B.** Correlations of the protein abundance values (log_10_-scaled) in the wild-type and deletion backgrounds across replicates. *r_s_*: Spearman’s rank correlation coefficient; *p*: p-value associated with the correlation. The robustness of the correlations was tested by five rounds of resampling.

**Figure S3.**
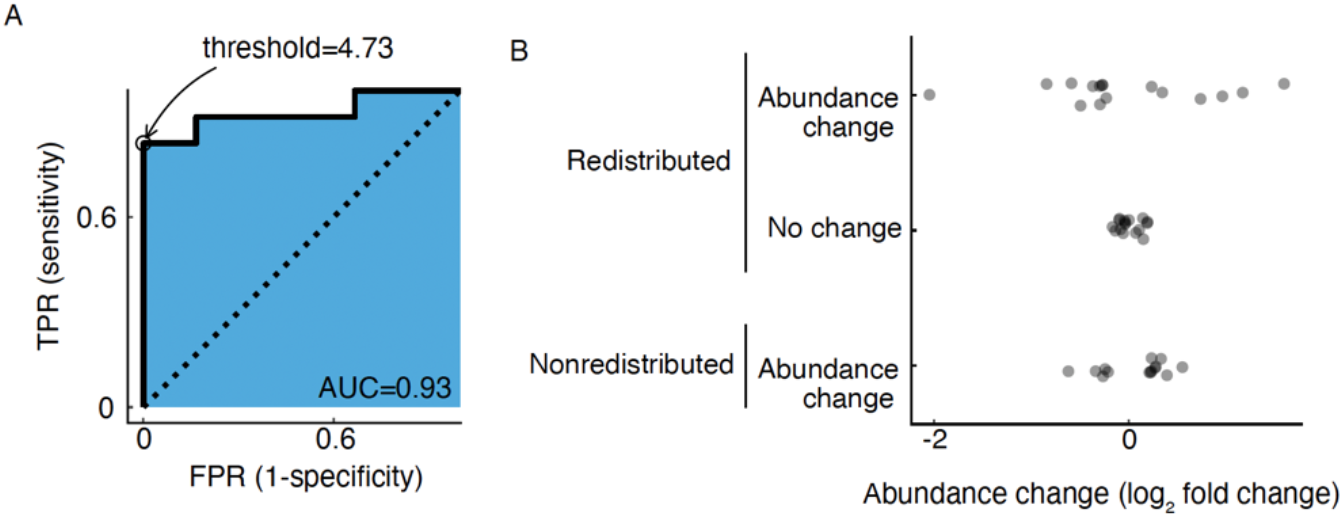
The classification of redistributed paralogs and comparison with the relative abundance change. **A.** Receiver Operating Characteristic (ROC) curve for the classification of the redistributed paralogs. **B.** Relative abundance change scores of paralogs stratified by the significance of redistribution and abundance change.

**Figure S4.**
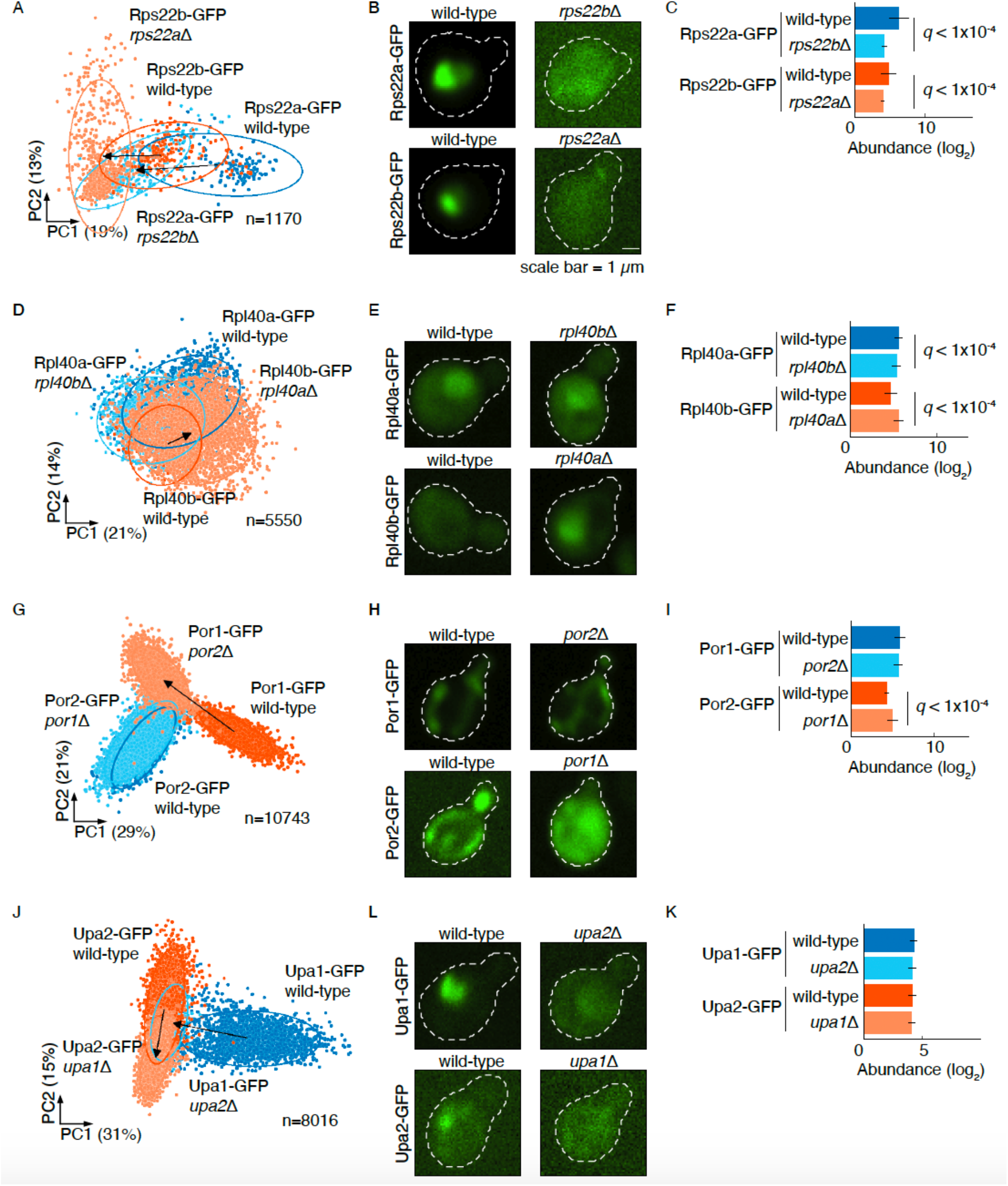
Redistribution, relative abundance and relocalization of paralog pairs. **A.** Redistribution of the Rps22a-Rps22b (in **A-C**), Rpl40a-Rpl40b (**D-F**), Por1-Por2 (**G-I**) and Upa1-Upa2 (**J-L**) pairs. Principal component analysis (PCA) of z-score normalized features is shown for each construct. For the redistributed paralog, the arrow connects the centroids of the clusters corresponding to the wild-type (dark blue/orange) and deletion (light blue/orange) backgrounds of the sister paralog. The percentage variances explained by the PCs are indicated in parentheses. **B.** Micrographs of representative yeast cells of respective paralog pairs. **C.** Relative abundance changes of the respective pairs. The error bars show the 95% confidence intervals of the means. *q*: FDR-corrected *p*-value. Wild-type (dark blue/orange) and deletion (light blue/orange) backgrounds of the paralog.

**Figure S5.**
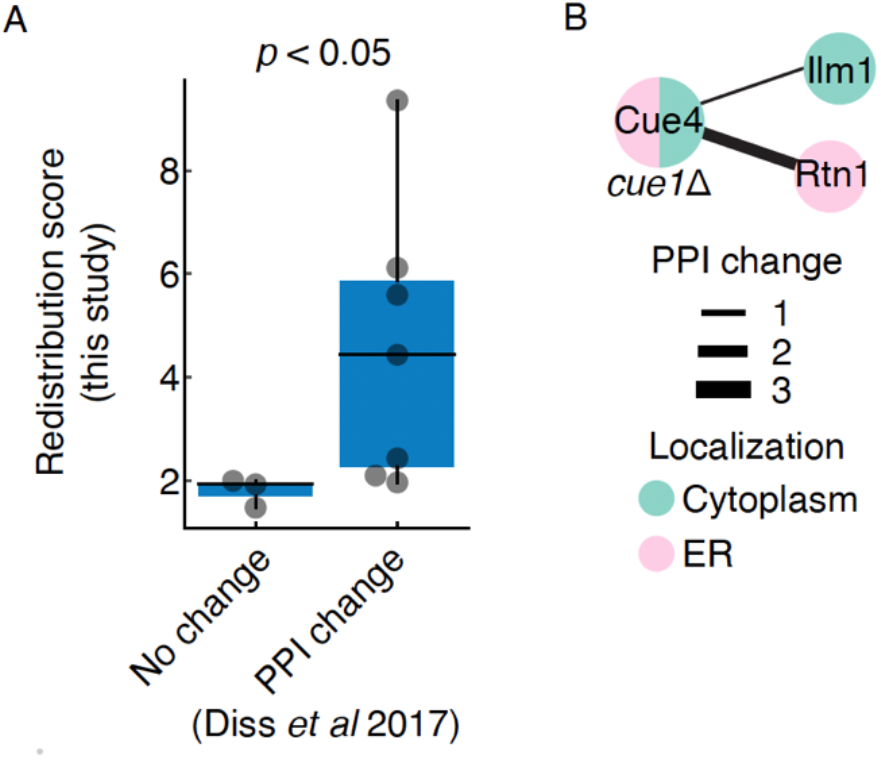
Redistributed paralogs show change in protein-protein interactions (PPIs). **A.** The comparison of the redistribution scores (this study) for proteins that show no change in PPIs with those that show at least one PPI change when their paralog is deleted. Statistical significance was determined using a two-sided Mann-Whitney U-test. **B.** Cue1-Cue4 represent an example of paralogous proteins in which only Cue4 shows redistribution (this study) and a corresponding PPI change upon *CUE1* deletion. The colors of the nodes indicate the subcellular compartments. Edges connect interacting proteins; edge thickness shows PPI change. PPI change was obtained from a previous study (Diss et al. 2017). In the boxplots, the central line indicates the median, the extent of the box is from the first quartile (Q1) to the third quartile (Q3), and the whiskers extend to Q1-1.5*IQR and Q3+1.5*IQR.

## Supplementary tables

Table S1: Yeast strains and plasmids used in this study.

Table S2: Manual inspection of paralog pairs and negative controls.

Table S3: Protein abundance

Table S4: The redistribution, relative abundance change and relocalization.

Table S5: Features of the paralogs.

## Supplementary data

Data S1: Features of single cells.

Data S2: The abundance per single cell.

## Notes

### Competing Interest Statement

The authors have declared no competing interest.

